# Fibroblasts as an Alternative to Mesenchymal Stem Cells with Successful Treatment and Immune Modulation in EAE Model of Multiple Sclerosis

**DOI:** 10.1101/2020.06.04.133249

**Authors:** Thomas E Ichim, Pete O’Heeron, Jesus Perez, Peter Liu, Wei-Ping Min, Santosh Kesari

## Abstract

The immune modulatory potential of mesenchymal stem cells (MSCs) is well known and is the basis for multiple clinical trials in treatment of autoimmune conditions. Unfortunately, MSCs are relatively rare, difficult to expand in culture, and methods of obtaining MSCs are complicated and expensive. In contrast, fibroblasts are found in copious amounts in various tissues, are a robust cellular population, and can be cultured without need for costs associated with culture media. Previous studies by our group and others have demonstrated fibroblasts possess regenerative activities. In the current study we demonstrated: a) fibroblasts inhibit mixed lymphocyte reaction; b) suppress T cell activation; c) inhibit DC maturation; and d) stimulate T regulatory (Treg) cell formation. Importantly, administration of fibroblasts in the experimental autoimmune encephalomyelitis (EAE) model of multiple sclerosis resulted in disease inhibition, which was abrogated upon depletion of Treg cells. This data, combined with existing clinical safety data on fibroblast administration, supports the clinical translation of fibroblast-based therapies for multiple sclerosis.

## Background

Multiple sclerosis (MS) is a T cell mediated autoimmune disorder targeting the myelin sheath which insulates neuronal bodies. The condition presents either as relapse remitting, which as the name suggests, undergoes periods of spontaneous remission, or primary progressive disease, in which the disease advances without remission. It is known that remission is associated with reduction of pathogenic T cells and upregulation of number and activity of T regulatory (Treg) cells (1), while progression is associate with augmentation of myelin-reactive T cells and suppression of Treg activity and number (2). Current treatments for MS include non-specific immune modulators such as Avonex, Capaxone, Rebif, Tysabri and Campath (3). These approaches possess various degrees of toxicity, adverse reactions associated with non-specific immune suppression, as well, display limited efficacy against primary progressive disease.

An ideal treatment for MS would possess the ability to suppress immunological aspects of the disease, while concurrently stimulating regeneration of injured neural tissues. Immunologically, it is believed that Treg cells play an important part of controlling disease progression. In addition to association between Treg and natural disease remission, studies have shown that efficacy of various clinically used treatments of MS associated with induction of Treg activity. Indeed, this has been shown in the case of interferon beta (4-10), as well as Copaxone (11-18), based interventions. Furthermore, animal studies have shown that adoptive administration of Treg cells in models of MS results in disease remissions (19-29). Alternatively, depletion of Tregs utilizing antibody or gene mediated approaches results in acceleration of the disease (30). Therefore, an ideal treatment for MS would possess ability to immune modulate through manipulation of the Treg compartment. Another aspect of MS is the neurodegeneration which occurs as a result of immunologically mediated demyelination. This demyelination results in progressive loss of neurons, which at the clinical level is observed as neurologically mediated deterioration. Chemical and other stimulators of endogenous neural stem cells have been shown to restore myelination in animal models, as well as reversing the pathology (31-52). This suggests it may be feasible to induce repair of tissues which have already been damaged by the immune system using agents which stimulate either oligodendrocyte function thereby promoting remyelination, as well as agents which stimulate endogenous neural stem cells, in order to regenerate neurons which have been damaged.

One approach which has been suggested to concurrently elicit regeneration while slowing down, or reversing immunological abnormalities in multiple sclerosis has been the utilization of cellular based therapies. While “stem cells” originally were believed to function therapeutically by restoring injured tissues, newer data demonstrated therapeutic activity correlates with generation of growth factors and/or antiapoptotic factors by stem cells. Indeed the mesenchymal stem cell type of therapies have been demonstrated to possess multiple immunological functions including: a) ability to suppress T cell activation (53-57); b) ability to suppress activation of NK cells (58-61); c) propensity to induce generation of Treg cells (62-64); and d) efficacy in a wide variety of models of autoimmune disease including the EAE model multiple sclerosis, autoimmune uveitis (65), and the collagen induce arthritis (CIA) model of rheumatoid arthritis (66). In addition to immune modulation, it has been demonstrated that MSCs possess the ability to regenerate neural tissue in models of MS, in part through stimulation of endogenous neural stem cells. Although MSCs possess multiple exiting properties, practical translation has been somewhat poor, with results obtained clinically only marginally effective. In addition, MSCs are difficult to isolate in bone marrow and adipose MSCs. They also require highly invasive extraction techniques.

Fibroblasts are a fundamental cell type in wound healing and have been previously demonstrated to possess some similarities to MSCs including CD73, CD90, and CD105 markers, as well as orthodox differentiation ability into bone, cartilage and adipose tissue. Although not studied in multiple sclerosis, a previous study explored whether fibroblasts isolated from skin may suppress the host immune response in a model of autoimmune arthritis. It was found that fibroblasts possessed the capacity to inhibit in vitro the proliferation of T lymphocytes. Fibroblasts also secrete modulatory molecules, such as prostaglandin E2 and nitric oxide, similar to MSCs. To assess their role in vivo, the collagen-induced arthritis model was used, and showed that similar to MSCs the intravenous injection of fibroblasts efficiently suppressed clinical signs of arthritis and delay of disease onset. This effect was associated with reduced inflammation as reflected by biological parameters and increased levels of IL-5, IL-10 and IL-13 in the spleens of treated mice (67). The CybroCell™ product is a clinical grade fibroblast population which has been demonstrated to possess preclinical and clinical efficacy in treatment of degenerative disc disease. In the current paper we assessed whether fibroblasts are superior immune modulators to MSCs, has well as assessed their therapeutic ability in an animal model of EAE.

## Materials and Methods

### Animal Model of Multiple Sclerosis

Adult male Lewis rats (9–10 weeks of age, 200–250 g) where assigned to four groups: normal control group (n = 12), BM-MSC group (n = 12), Adipose MSC group (n = 12), and Fibroblast group (n=12). Experimental autoimmune encephalomyelitis (EAE) was induced by subcutaneous injection of guinea pig spinal cord homogenate (GPSCH) emulsified at a 1:1 ratio with complete Freund adjuvant (CFA) containing heat killed Mycobacterium tuberculosis. Each rat received an intraperitoneal injection of 300 ng Pertussis toxin (Sigma-Aldrich, St. Louis, MO, USA) in 0.1 ml distilled water immediately after the subcutaneous injection and again 48hours later. Cells where injected at a concentration of 1 million cells per rat intravenously subsequent to administration of GPSCH. The clinical manifestations of EAE were assessed daily until the time of sacrifice. Disease severity was scored on a 5-point scale: 0 = no signs, 1 = partial loss of tail tonicity, 2 = loss of tail tonicity, 3 = unsteady gait and mild paralysis, 4 = hind limb paralysis and incontinence, and 5 = moribund or death. Disease scoring was performed by pathologists blinded to treatment conditions.

### Antibody Depletion and Cytokine Assessments

Depletion of Treg cells was performed using goat anti-rat CD25 antibody by administration of the antibody every second day intraperitoneally. Confirmation of Treg depletion was performed by flow cytometric assessment. When >75% depletion of CD25 cells at 7 days post EAE induction was accomplished (confirmed by different epitope targeting antibody), animals where considered “depleted”. For assessment of cytokine levels, blood was drawn from the tail vein and IL-10 and IL-17 levels where assessed from plasma using Enzyme Linked Immunosorbent Assay (ELISA). Assessments where made at days 0, 18, and 31.

### Fibroblasts

Rat dermal fibroblasts where obtained from ATCC and propagated in Optimem media using 10% FCS and pen-strep in fully humidified atmosphere. In some experiments human fibroblasts where utilized and grown under similar conditions.

### Mixed Lymphocyte Reactions

Dermal fibroblasts where obtained from ATCC and maintained in DMEM media with 10% FCS in a fully humidified environment, with penicillin/streptomycin mixture and non-essential amino acids. Cells where harvested at 75% confluence by trypsinization and plated with immature dendritic cells at day 5 of DC maturation. Cells where plated in 12 well plates with 100,000 fibroblasts per 1,000,000 DC, 500,000 fibroblasts per 1,000,000 DC and 1,000,000 fibroblasts per 1,000,000 DC. After 48 hours of culture cells where extracted and CD40 expression was assessed by flow cytometry.

Generation of DC was performed by culturing monocytes in GM-CSF and IL-4 for 5 days according to the method of Inaba et al. and subsequently matured by addition of TNF-alpha on day 5 before the coculture. In some experiments LPS was added to the fibroblasts at a concentration of 5 ug/ml.

## Results

### Reduction of Multiple Sclerosis Pathology in the Experimental Allergic Encephalomyelitis (EAE) Model by Fibroblasts is Superior to Mesenchymal Stem Cells

Previous studies have demonstrated fibroblasts possess various similarities to mesenchymal stem cells (MSCs) including surface markers, morphology, and some aspects of immune modulation (68). In order to compare therapeutic activity between fibroblasts and MSCs, we utilized the rat model of multiple sclerosis termed experimental allergic encephalomyelitis (EAE). This model consists of immunization with xenogeneic spinal cord homogenate as a source of myelin basic protein (MBP). Also, utilizing a strong adjuvant to break self-tolerance, it combines with pertussis toxin to permeabilize the blood brain barrier. For comparison, MSCs where utilized from adipose and bone marrow sources.

All cells where administered at a concentration of 1 million cells, intravenously, the same day as administration of xenogeneic spinal cord homogenate. As seen in Figure 1, BM-MSCs possessed the least EAE inhibitory activity, while adipose-MSCs possessed slightly stronger inhibitory activity. Fibroblasts induced potent reduction of disease progression, with no clinical symptomology on day 24, in contrast, both BM and adipose derived MSCs treated groups had pathology through completion of experiments at day 31.

**Figure 1:**
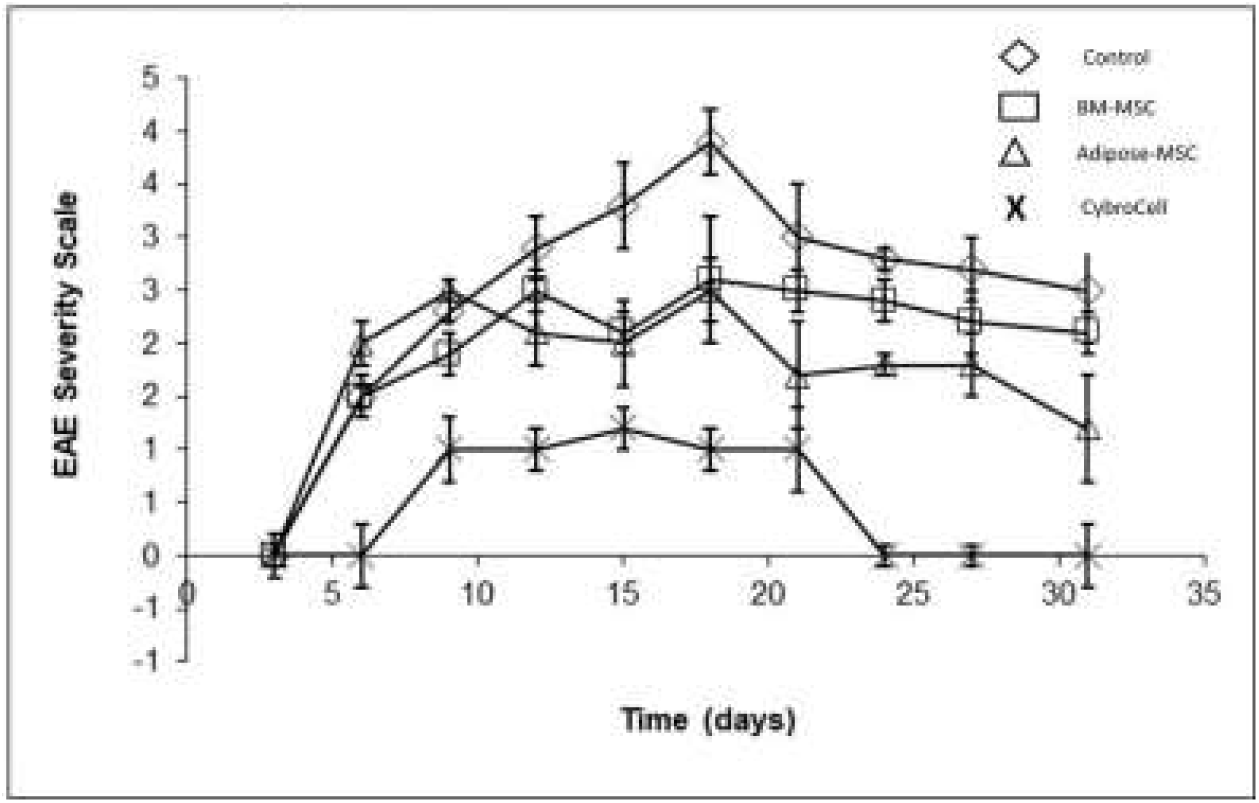
Superior Inhibition of EAE Pathology by Fibroblasts Compared to MSC.

### In Vitro Mechanisms of Fibroblast Immune Modulatory Activity

While previous studies showed immune modulatory activities of fibroblasts, including ability to suppress mixed lymphocyte reaction, direct suppression of autoantigen specific responses has not been demonstrated to our knowledge. This is fundamentally important due to the qualitative differences between an alloantigen-specific immune reaction, in which numerous T cell clones are responding, compared to antigen-specific reactions in which only one or a number of T cell clones are expanding. Accordingly, T cells specific to myelin basic protein were generated by in vitro stimulation with antigen presenting cells and interleukin-2 administration. An increasing number of fibroblasts were shown to dose-dependently inhibit proliferation of antigen specific reactive T cells (Figure 2a). Additionally, utilizing a similar system, a question was contemplated whether fibroblasts can elicit generation of T regulatory cells (Treg). It is known that Treg cells are capable of suppressing autoimmunity in a variety of situations, leading to the question of whether fibroblasts are capable of inducing generation of this cell type. As seen in Figure 2b, increasing numbers of FoxP3 cells where observed in MPB-reacting T cells when fibroblasts where added.

**Figure 2:**
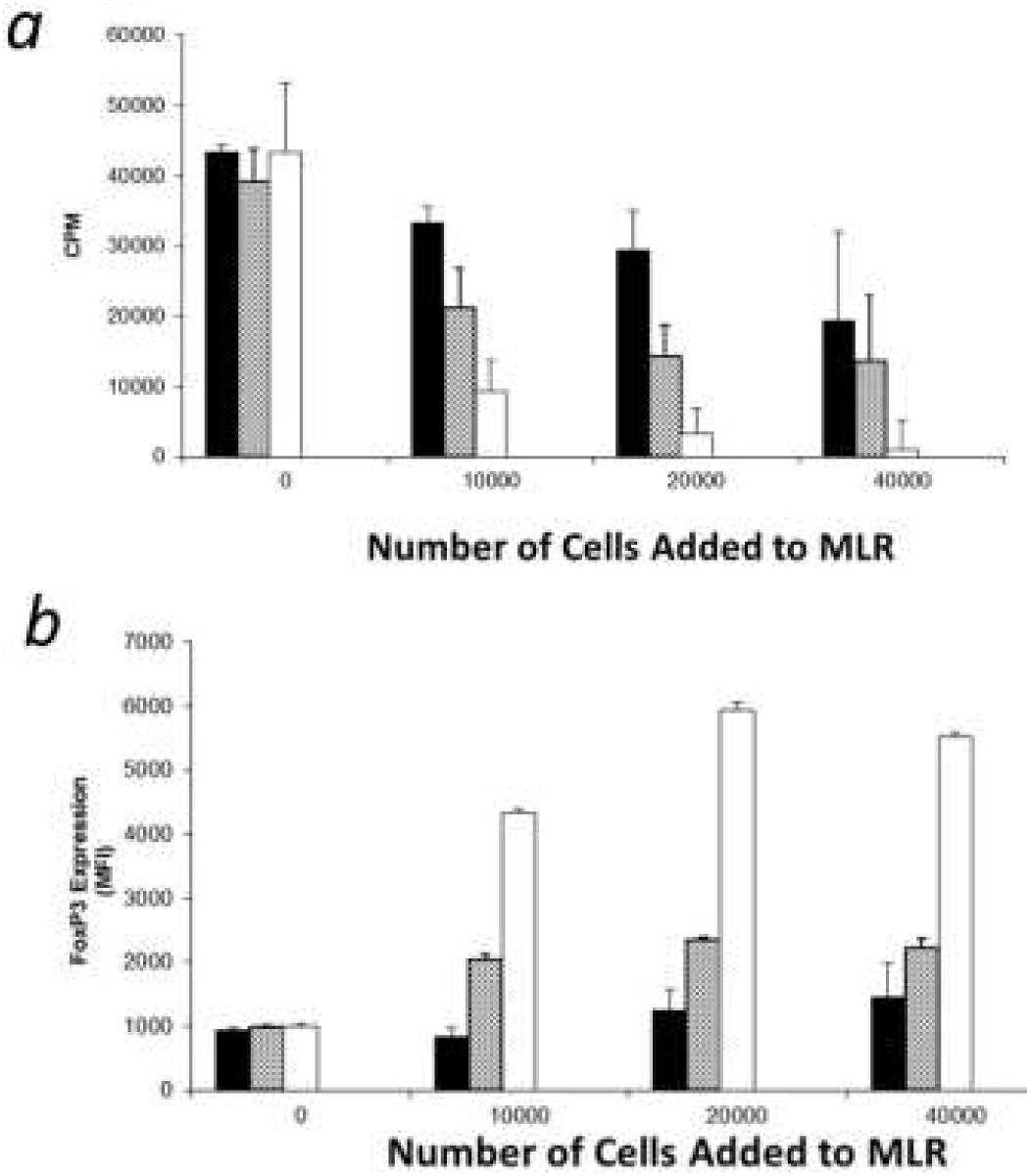
Fibroblasts Modulate Antigen-Specific T cell Responses.

### Fibroblasts Block IL-17 Inflammatory Pathway

It is known Treg cells are reciprocal cells to Th17 cells, and it is also known the Th17 CD4 T cell subsets plays a major role in pathology of EAE as well as numerous other autoimmune disorders in man and mice (69-79). Accordingly, we assessed the IL-17 mediated pathway in vitro and in vivo. Fibroblasts dose dependently suppressed macrophage production of TNF-alpha subsequent to toll-like receptor (TLR-4) stimulation with lipopolysaccharide (Figure 3a). Given that IL-17 is a hallmark of pathogenic Th17 cells, which are upstream of TNF-alpha producing cells in the immunological cascade, we assessed in vitro whether IL-17 was reduced by fibroblasts in response to IL-6 induction. We observed a dose dependent inhibition of IL-17 production in vitro (Figure 3b). In order to assess in vivo relevance of these findings, we sought to determine levels of IL-17, as well as its reciprocal cytokine IL-10. This is made by Treg cells and is known to activate Treg cells by assessing levels of these cytokines from the peripheral blood of rats undergoing EAE at days 0, 18 and 31. Quantification of cytokines was performed in rats who received control saline, fibroblasts, BM-MSCs, and adipose MSCs. Similar changes were observed in vivo with suppression of IL-17 and stimulation of Treg cells (Figure 3c and 3d).

**Figure 3:**
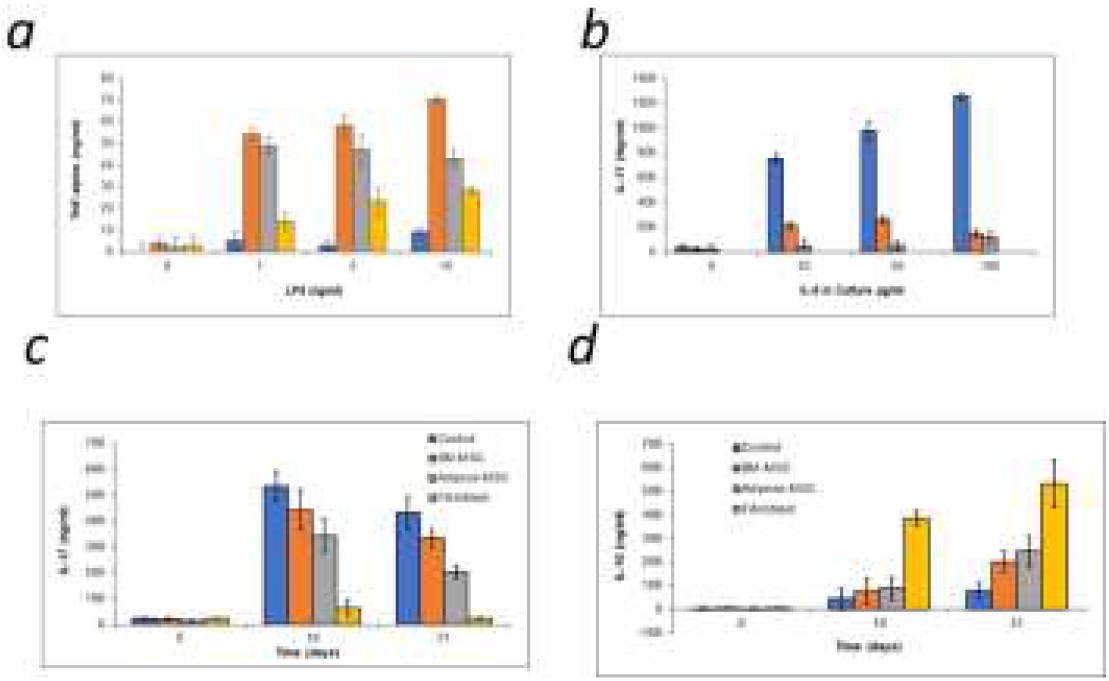
Fibroblasts Block IL-17 Pathway in Vitro and in Vivo.

### Inhibition of Dendritic Cell Maturation by Fibroblasts

The fundamental importance of dendritic cells (DCs) cannot be overstated in both activation and suppression of immune responses. Mature DCs are uniquely capable of activating naïve T cells through their high and dense expression of membrane-bound costimulatory molecules such as CD40, CD80, and CD86 and the soluble costimulatory signal IL-12. Immature dendritic cells possess ability to induce immunological tolerance through their high levels of co-inhibitory molecules such as IL-10 and PD-L1 (80-94). Given that fibroblasts are associated with post-injury healing, and that upregulation of immune inhibitory molecules is known to occur in the healing phase of injury, we sought to investigate whether DC maturation was altered by fibroblasts using a co-culture system. Suppression of TNF/LPS induced DC maturation was observed in terms of downregulation of CD40, CD80, CD86, and IL-12, along with upregulation of the inhibitory molecules IL-10, IL-1RA, and PD-L1 (Figure 4a-g). In accordance with the in vivo data demonstrating superior activity of fibroblasts to MSCs, fibroblasts where superior to bone BM and adipose derived MSCs at suppressing DC maturation (Figure 5a-g).

**Figure 4:**
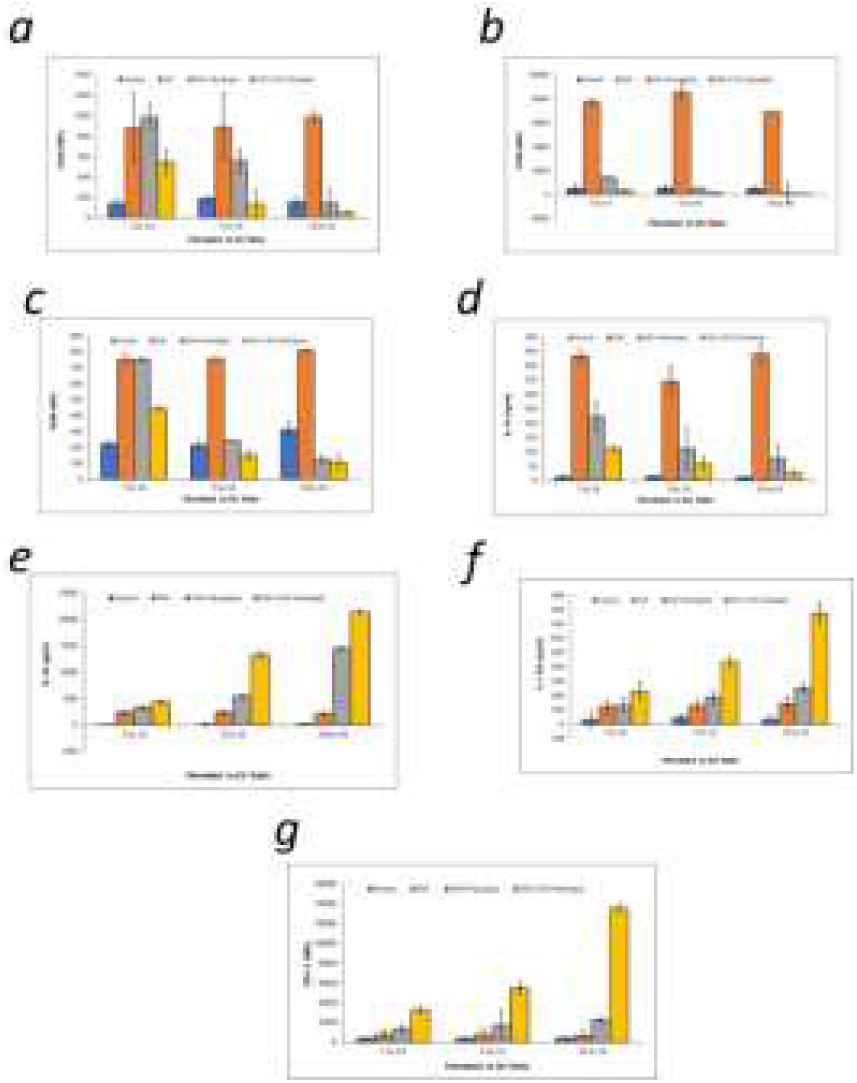
Fibroblasts Inhibit DC Maturation.

**Figure 5:**
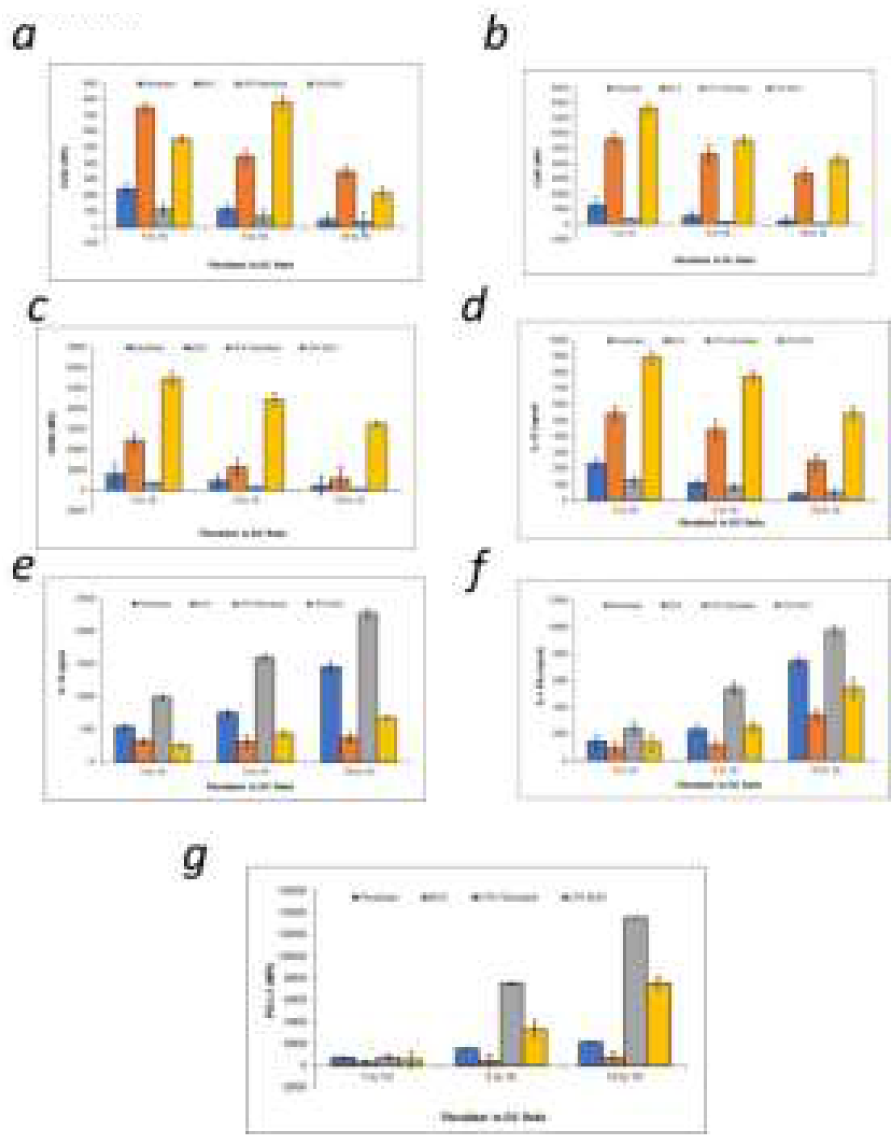
Fibroblasts are Superior to MSC at inhibiting DC Maturation.

### Depletion of Treg Results in Abrogation of Protection from EAE Pathology by Fibroblasts

Given the findings that fibroblasts reduce generation of IL-17 cytokine in response to IL-6 in vitro and inhibition IL-17 in vivo while stimulating IL-10, the potential of fibroblasts inhibiting EAE through a Treg dependent manner was considered. This possibility is further strengthened by our findings that fibroblasts stimulate generation of Treg in vitro. Accordingly, a series of experiments was conducted in order to determine whether antibody mediated depletion of Treg would alter efficacy of fibroblasts at reduction of EAE pathology. As seen in Figure 6, reduction of EAE pathology in response to fibroblast administration was significantly abrogated in animals in which Treg where depleted but not in isotype control immunized animals.

**Figure 6:**
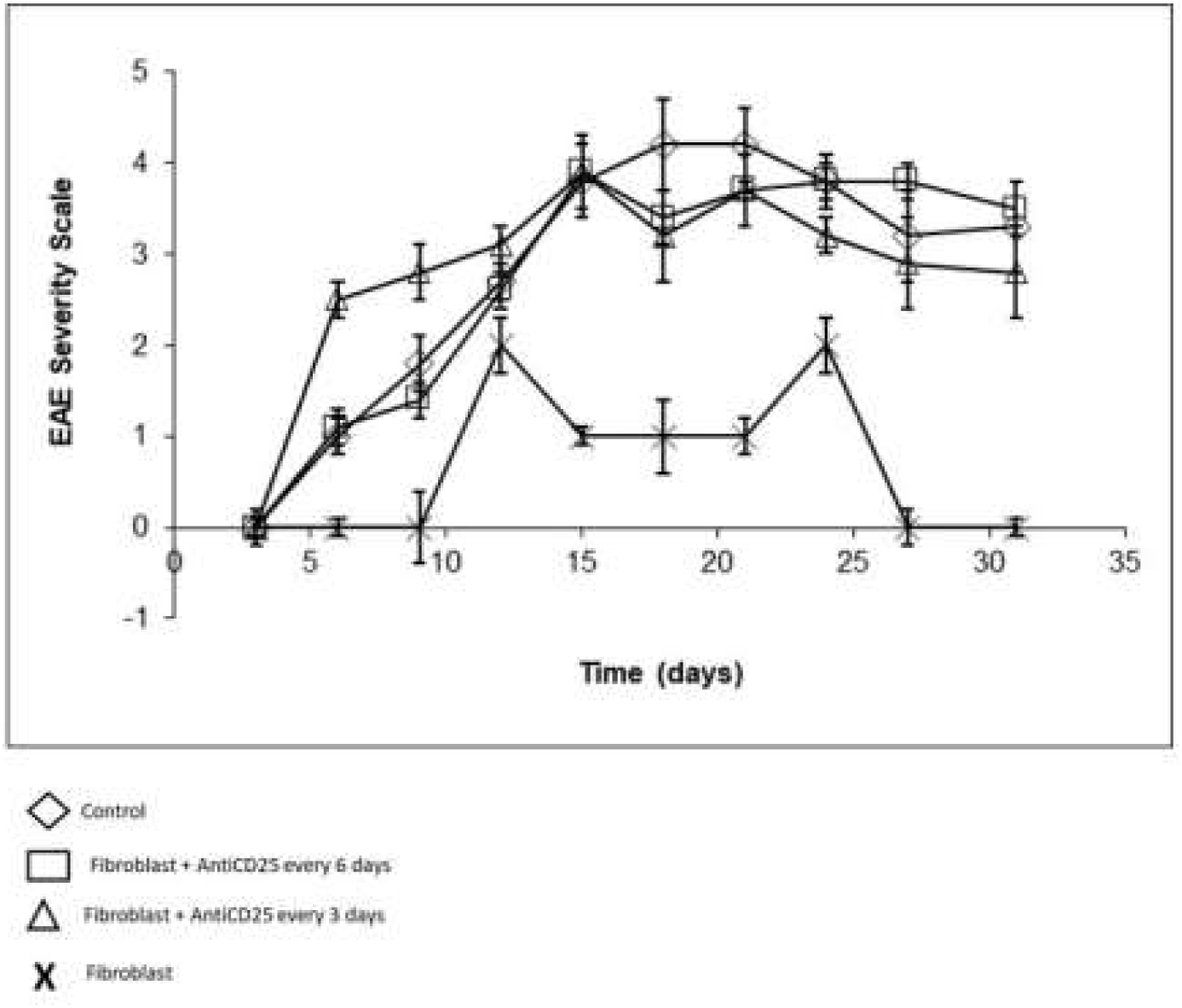
Depletion of Treg Abrogates Therapeutic Effect of Fibroblasts on EAE Pathology.

### Regenerative Effects of Fibroblasts on EAE Pathology

Although numerous immune modulatory agents have been explored in the context of EAE as a preclinical model for translational development, relatively little work has been performed in examining the possibility of agents which stimulate regeneration of tissue subsequent to injury. Accordingly, we examined the ability of neuronal remyelination by assessing whether the number of remyelinated neurons increased subsequent to treatment. As seen in Figure 7a, an increase in remyelination was observed, which was more profound with fibroblasts as compared to BM and adipose derived MSCs. Additionally, slices of the dentate gyrus revealed cells possessing the proliferating cell nuclear antigen (PCNA), which is a marker of cellular proliferation (Figure 7b). These data suggest that fibroblasts reduce pathology of EAE not only by stimulation of inhibitory immune responses, primarily such as Treg and immature/tolerogenic DC, but also fibroblasts mediate a regenerative effect.

**Figure 7:**
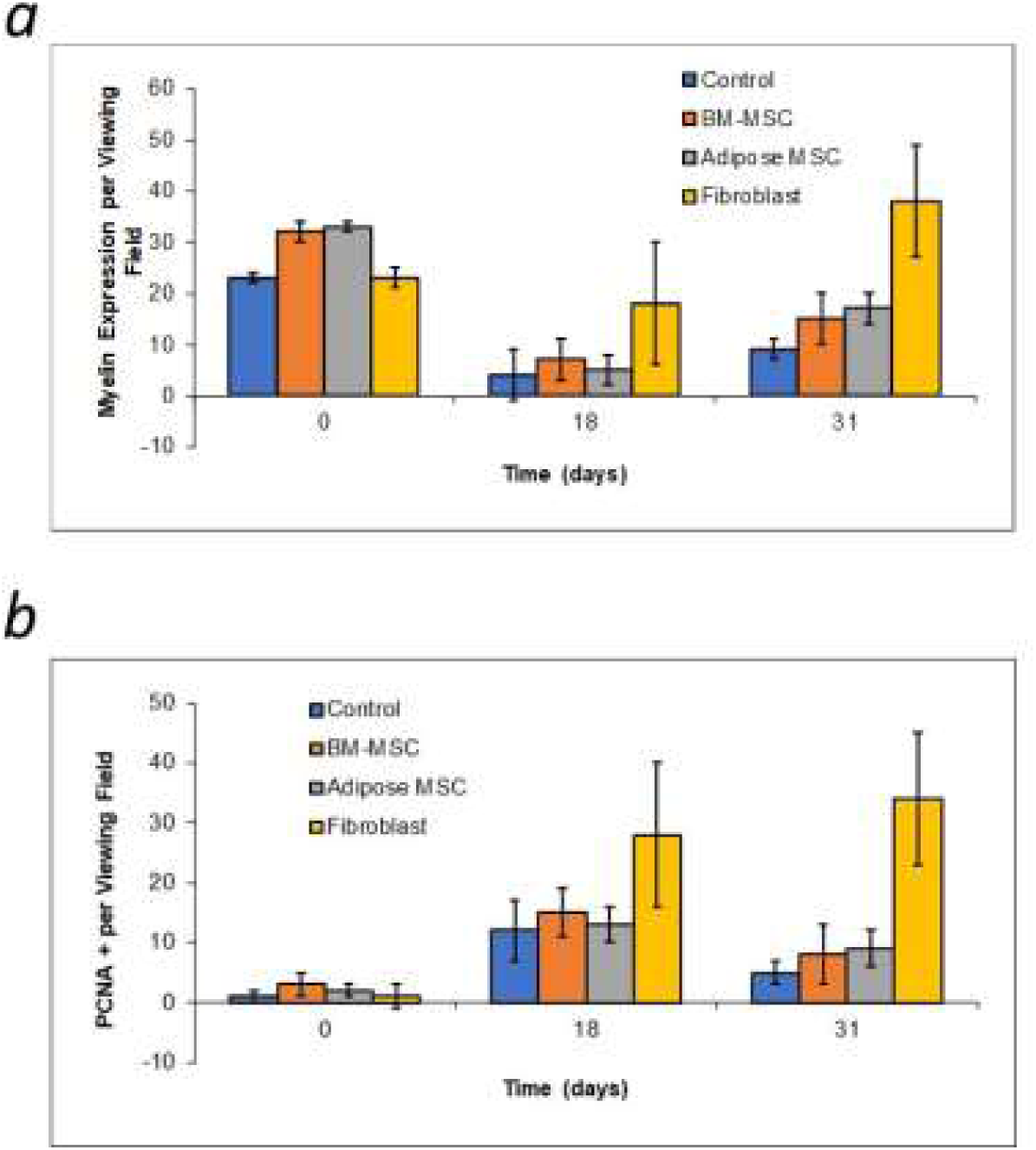
Regenerative Effects of Fibroblasts in EAE Model.

## Discussion

Regenerative therapy for MS has historically been limited to stem cell approaches. Autologous hematopoietic stem cell use post-immune ablation has demonstrated some level of efficacy, however cost and potential adverse reactions of full-immune compromise make this treatment difficult to implement on a large scale. The use of MSCs has demonstrated promise in animal models, but, clinical responses have been mediocre. The cost of culturing MSCs, as well as difficulties in acquisition and quality control represents hurdles in the commercialization of these cells. In the current paper we demonstrate superior efficacy of fibroblasts to MSCs in terms of reducing progression of disease pathology, stimulating Treg formation, inhibiting IL-17 and augmenting IL-10.

Utilization of fibroblasts as a substitute for MSCs has been previously proposed by our group based on similar phenotypic and morphological characteristics between the two cell types. Previously published data supporting that fibroblasts, similar to MSCs, possess ability to differentiate along the orthodox pathway, such as chondrocytes, adipocytes, and osteocytes. In addition to studies, which demonstrated efficacy of fibroblasts for differentiating into chondrocytes in vivo and generating improvement in animal models of degenerative disc disease, an independent group reported similar findings. This strongly supports development of fibroblast-based products as an alternate to MSCs. The recent FDA IND clearance to initiate clinical trials using fibroblasts for treatment of patients with degenerative disc disease, is further support for this. The demonstration of superior inhibition of multiple sclerosis-like pathology using fibroblasts as compared to MSCs cannot be accounted for by different passage numbers since these were equalized between groups. Furthermore, an increased efficacy using adipose derived MSCs compared to bone marrow was shown. Previous studies which have suggested superior immune modulatory effects of adipose MSCs compared to BM MSCs. While fibroblasts used in the experiments were dermal derived, the possible optimization of therapeutic activity by choosing fibroblasts from other sources is an intriguing question which is currently under investigation.

Mechanistically, the suppression of IL-17 production in vitro and in vivo appeared to be related to efficacy of fibroblasts. Other studies have suggested that MSCs possess ability to suppress IL-17, however, in these experiments fibroblasts where markedly superior. Some studies have shown wound healing is associated with potent immune modulation, accordingly, given the potent role of fibroblasts in wound healing, cytokine mediated immune modulation activity of fibroblasts appears to be quite potent. Other soluble factors have not been examined but are the topic of current investigation. One particular area warranting further examination is whether fibroblasts are mediating therapeutic effects by secretion of exosomes. Our preliminary data supports the ability of exosomes to modulate some aspects of the regenerative process. Indeed, future studies may seek to optimize therapy by administration of exosomes together with cellular therapy.

It has previously been demonstrated by us (WPM and TEI) that immature DC give rise to tolerogenic DC which in turn cause activation of more Treg. This “inhibitory feedback loop” demonstrated to be responsible for maintaining allograft tolerance in the B6 to BALB/c cardiac heterotopic transplant model. Accordingly, it may be feasible to imagine that fibroblasts are giving rise to immature, or “tolerogenic” DC, which in turn cause generation of Treg cells. Indeed, the data described supports the fundamental role of Treg in fibroblast-mediated reduction for EAE pathology resulting in depletion of the cell loss of therapeutic activity. Future studies to combine Treg cells with fibroblast administration and/or tolerogenic DC are underway. This gives rise to theconcept that the fibroblast may not only serve as a monotherapy for treatment of MS, but may also increase efficacy of existing cell therapies for this condition. Furthermore, certain drugs on the market are believed to function, intra alia, through stimulation of Treg cells. The combination of fibroblasts with drugs such as copaxone and interferon beta are being investigated.

Endogenous neural stem cells have been reported to accelerate repair of various neurological injuries including stroke, traumatic brain injury, and EAE. The possibility of stimulating endogenous neural stem cells using external approaches has been investigated with some success using approaches such as hCG administration. Recently administration of MSCs has been shown to enhance proliferation of endogenous neural stem cells and may have been a mechanism of action for some types of cellular therapy. The current study demonstrated fibroblasts can enhance neural stem cells, which offers the possibility of repairing injured tissue. Additionally, our data demonstrated augmentation of myelination subsequent to treatment. Whether this is result of de novo regeneration of oligodendrocytes from progenitor cells, or whether activation of existing oligodendrocyte cells is still under investigation. This data is particularly interesting in light of a publication by Nessler et al. who showed no effect of MSCs on cuprizone induced demyelination, which is non-immune mediated (95).

In conclusion, we describe the novel ability of fibroblasts to reduce autoimmunity in the EAE model of multiple sclerosis. The possibility of utilizing less expensive fibroblasts as a superior type over MSCs for therapeutic use is will lead to further development and testing.

## References

1. Rodi, M., Dimisianos, N., de Lastic, A. L., Sakellaraki, P., Deraos, G., Matsoukas, J., Papathanasopoulos, P., and Mouzaki, A. (2016) Regulatory Cell Populations in Relapsing-Remitting Multiple Sclerosis (RRMS) Patients: Effect of Disease Activity and Treatment Regimens. Int J Mol Sci 17

2. Elong Ngono, A., Pettre, S., Salou, M., Bahbouhi, B., Soulillou, J. P., Brouard, S., and Laplaud, D. A. (2012) Frequency of circulating autoreactive T cells committed to myelin determinants in relapsing-remitting multiple sclerosis patients. Clin Immunol 144, 117–126

3. Saez-Torres, I., Brieva, L., Espejo, C., Barrau, M. A., Montalban, X., and Martinez-Caceres, E. M. (2002) Specific proliferation towards myelin antigens in patients with multiple sclerosis during a relapse. Autoimmunity 35, 45–50

4. Ebrahimimonfared, M., Ganji, A., Zahedi, S., Nourbakhsh, P., Ghasami, K., and Mosayebi, G. (2018) Characterization of Regulatory T-Cells in Multiple Sclerosis Patients Treated with Interferon Beta-1a. CNS Neurol Disord Drug Targets 17, 113–118

5. Wang, D., Ghosh, D., Islam, S. M., Moorman, C. D., Thomason, A. E., Wilkinson, D. S., and Mannie, M. D. (2016) IFN-beta Facilitates Neuroantigen-Dependent Induction of CD25+ FOXP3+ Regulatory T Cells That Suppress Experimental Autoimmune Encephalomyelitis. J Immunol 197, 2992–3007

6. Cosentino, M., Zaffaroni, M., Trojano, M., Giorelli, M., Pica, C., Rasini, E., Bombelli, R., Ferrari, M., Ghezzi, A., Comi, G., Livrea, P., Lecchini, S., and Marino, F. (2012) Dopaminergic modulation of CD4+CD25(high) regulatory T lymphocytes in multiple sclerosis patients during interferon-beta therapy. Neuroimmunomodulation 19, 283–292

7. Chen, M., Chen, G., Deng, S., Liu, X., Hutton, G. J., and Hong, J. (2012) IFN-beta induces the proliferation of CD4+CD25+Foxp3+ regulatory T cells through upregulation of GITRL on dendritic cells in the treatment of multiple sclerosis. J Neuroimmunol 242, 39–46

8. Kieseier, B. C. (2011) The mechanism of action of interferon-beta in relapsing multiple sclerosis. CNS Drugs 25, 491–502

9. Vandenbark, A. A., Huan, J., Agotsch, M., La Tocha, D., Goelz, S., Offner, H., Lanker, S., and Bourdette, D. (2009) Interferon-beta-1a treatment increases CD56bright natural killer cells and CD4+CD25+ Foxp3 expression in subjects with multiple sclerosis. J Neuroimmunol 215, 125–128

10. Penton-Rol, G., Cervantes-Llanos, M., Cabrera-Gomez, J. A., Alonso-Ramirez, R., Valenzuela-Silva, C., Rodriguez-Lara, R., Montero-Casimiro, E., Bello-Rivero, I., and Lopez-Saura, P. (2008) Treatment with type I interferons induces a regulatory T cell subset in peripheral blood mononuclear cells from multiple sclerosis patients. Int Immunopharmacol 8, 881–886

11. Hong, J., Li, N., Zhang, X., Zheng, B., and Zhang, J. Z. (2005) Induction of CD4+CD25+ regulatory T cells by copolymer-I through activation of transcription factor Foxp3. Proc Natl Acad Sci U S A 102, 6449–6454

12. Horani, A., Muhanna, N., Pappo, O., Melhem, A., Alvarez, C. E., Doron, S., Wehbi, W., Dimitrios, K., Friedman, S. L., and Safadi, R. (2007) Beneficial effect of glatiramer acetate (Copaxone) on immune modulation of experimental hepatic fibrosis. Am J Physiol Gastrointest Liver Physiol 292, G628–638

13. Jee, Y., Piao, W. H., Liu, R., Bai, X. F., Rhodes, S., Rodebaugh, R., Campagnolo, D. I., Shi, F. D., and Vollmer, T. L. (2007) CD4(+)CD25(+) regulatory T cells contribute to the therapeutic effects of glatiramer acetate in experimental autoimmune encephalomyelitis. Clin Immunol 125, 34–42

14. Weber, M. S., Hohlfeld, R., and Zamvil, S. S. (2007) Mechanism of action of glatiramer acetate in treatment of multiple sclerosis. Neurotherapeutics 4, 647–653

15. Saresella, M., Marventano, I., Longhi, R., Lissoni, F., Trabattoni, D., Mendozzi, L., Caputo, D., and Clerici, M. (2008) CD4+CD25+FoxP3+PD1-regulatory T cells in acute and stable relapsing-remitting multiple sclerosis and their modulation by therapy. FASEB J 22, 3500–3508

16. Cui, G., Zhang, Y., Gong, Z., Zhang, J. Z., and Zang, Y. Q. (2009) Induction of CD4+CD25+Foxp3+ regulatory T cell response by glatiramer acetate in type 1 diabetes. Cell Res 19, 574–583

17. Haas, J., Korporal, M., Balint, B., Fritzsching, B., Schwarz, A., and Wildemann, B. (2009) Glatiramer acetate improves regulatory T-cell function by expansion of naive CD4(+)CD25(+)FOXP3(+)CD31(+) T-cells in patients with multiple sclerosis. J Neuroimmunol 216, 113–117

18. Lalive, P. H., Neuhaus, O., Benkhoucha, M., Burger, D., Hohlfeld, R., Zamvil, S. S., and Weber, M. S. (2011) Glatiramer acetate in the treatment of multiple sclerosis: emerging concepts regarding its mechanism of action. CNS Drugs 25, 401–414

19. Duplan, V., Beriou, G., Heslan, J. M., Bruand, C., Dutartre, P., Mars, L. T., Liblau, R. S., Cuturi, M. C., and Saoudi, A. (2006) LF 15-0195 treatment protects against central nervous system autoimmunity by favoring the development of Foxp3-expressing regulatory CD4 T cells. J Immunol 176, 839–847

20. Wang, Z., Hong, J., Sun, W., Xu, G., Li, N., Chen, X., Liu, A., Xu, L., Sun, B., and Zhang, J. Z. (2006) Role of IFN-gamma in induction of Foxp3 and conversion of CD4+ CD25-T cells to CD4+ Tregs. J Clin Invest 116, 2434–2441

21. Ochoa-Reparaz, J., Riccardi, C., Rynda, A., Jun, S., Callis, G., and Pascual, D. W. (2007) Regulatory T cell vaccination without autoantigen protects against experimental autoimmune encephalomyelitis. J Immunol 178, 1791–1799

22. Stephens, L. A., Malpass, K. H., and Anderton, S. M. (2009) Curing CNS autoimmune disease with myelin-reactive Foxp3+ Treg. Eur J Immunol 39, 1108–1117

23. Benkhoucha, M., Santiago-Raber, M. L., Schneiter, G., Chofflon, M., Funakoshi, H., Nakamura, T., and Lalive, P. H. (2010) Hepatocyte growth factor inhibits CNS autoimmunity by inducing tolerogenic dendritic cells and CD25+Foxp3+ regulatory T cells. Proc Natl Acad Sci U S A 107, 6424–6429

24. Matsushita, T., Horikawa, M., Iwata, Y., and Tedder, T. F. (2010) Regulatory B cells (B10 cells) and regulatory T cells have independent roles in controlling experimental autoimmune encephalomyelitis initiation and late-phase immunopathogenesis. J Immunol 185, 2240–2252

25. Rynda-Apple, A., Huarte, E., Maddaloni, M., Callis, G., Skyberg, J. A., and Pascual, D. W. (2011) Active immunization using a single dose immunotherapeutic abates established EAE via IL-10 and regulatory T cells. Eur J Immunol 41, 313–323

26. Barsheshet, Y., Wildbaum, G., Levy, E., Vitenshtein, A., Akinseye, C., Griggs, J., Lira, S. A., and Karin, N. (2017) CCR8(+)FOXp3(+) Treg cells as master drivers of immune regulation. Proc Natl Acad Sci U S A 114, 6086–6091

27. Zilkha-Falb, R., Gurevich, M., and Achiron, A. (2017) Experimental Autoimmune Encephalomyelitis Ameliorated by Passive Transfer of Polymerase 1-Silenced MOG35-55 Lymphatic Node Cells: Verification of a Novel Therapeutic Approach in Multiple Sclerosis. Neuromolecular Med 19, 406–412

28. Kim, B. S., Lu, H., Ichiyama, K., Chen, X., Zhang, Y. B., Mistry, N. A., Tanaka, K., Lee, Y. H., Nurieva, R., Zhang, L., Yang, X., Chung, Y., Jin, W., Chang, S. H., and Dong, C. (2017) Generation of RORgammat(+) Antigen-Specific T Regulatory 17 Cells from Foxp3(+) Precursors in Autoimmunity. Cell Rep 21, 195–207

29. Kim, Y. C., Zhang, A. H., Yoon, J., Culp, W. E., Lees, J. R., Wucherpfennig, K. W., and Scott, D. W. (2018) Engineered MBP-specific human Tregs ameliorate MOG-induced EAE through IL-2-triggered inhibition of effector T cells. J Autoimmun 92, 77–86

30. Ghosh, D., Curtis, A. D., 2nd, Wilkinson, D. S., and Mannie, M. D. (2016) Depletion of CD4+ CD25+ regulatory T cells confers susceptibility to experimental autoimmune encephalomyelitis (EAE) in GM-CSF-deficient Csf2-/-mice. J Leukoc Biol 100, 747–760

31. Picard-Riera, N., Decker, L., Delarasse, C., Goude, K., Nait-Oumesmar, B., Liblau, R., Pham-Dinh, D., and Baron-Van Evercooren, A. (2002) Experimental autoimmune encephalomyelitis mobilizes neural progenitors from the subventricular zone to undergo oligodendrogenesis in adult mice. Proc Natl Acad Sci U S A 99, 13211–13216

32. Magalon, K., Cantarella, C., Monti, G., Cayre, M., and Durbec, P. (2007) Enriched environment promotes adult neural progenitor cell mobilization in mouse demyelination models. Eur J Neurosci 25, 761–771

33. Aharoni, R., Aizman, E., Fuchs, O., Arnon, R., Yaffe, D., and Sarig, R. (2009) Transplanted myogenic progenitor cells express neuronal markers in the CNS and ameliorate disease in experimental autoimmune encephalomyelitis. J Neuroimmunol 215, 73–83

34. Gordon, D., Pavlovska, G., Uney, J. B., Wraith, D. C., and Scolding, N. J. (2010) Human mesenchymal stem cells infiltrate the spinal cord, reduce demyelination, and localize to white matter lesions in experimental autoimmune encephalomyelitis. J Neuropathol Exp Neurol 69, 1087–1095

35. Banisadr, G., Frederick, T. J., Freitag, C., Ren, D., Jung, H., Miller, S. D., and Miller, R. J. (2011) The role of CXCR4 signaling in the migration of transplanted oligodendrocyte progenitors into the cerebral white matter. Neurobiol Dis 44, 19–27

36. Pedersen, D. S., Fredericia, P. M., Pedersen, M. O., Stoltenberg, M., Penkowa, M., Danscher, G., Rungby, J., and Larsen, A. (2012) Metallic gold slows disease progression, reduces cell death and induces astrogliosis while simultaneously increasing stem cell responses in an EAE rat model of multiple sclerosis. Histochem Cell Biol 138, 787–802

37. Khezri, S., Javan, M., Goudarzvand, M., Semnanian, S., and Baharvand, H. (2013) Dibutyryl cyclic AMP inhibits the progression of experimental autoimmune encephalomyelitis and potentiates recruitment of endogenous neural stem cells. J Mol Neurosci 51, 298–306

38. Grade, S., Bernardino, L., and Malva, J. O. (2013) Oligodendrogenesis from neural stem cells: perspectives for remyelinating strategies. Int J Dev Neurosci 31, 692–700

39. Pazhoohan, S., Satarian, L., Asghari, A. A., Salimi, M., Kiani, S., Mani, A. R., and Javan, M. (2014) Valproic Acid attenuates disease symptoms and increases endogenous myelin repair by recruiting neural stem cells and oligodendrocyte progenitors in experimental autoimmune encephalomyelitis. Neurodegener Dis 13, 45–52

40. Mecha, M., Feliu, A., Carrillo-Salinas, F. J., Mestre, L., and Guaza, C. (2013) Mobilization of progenitors in the subventricular zone to undergo oligodendrogenesis in the Theiler’s virus model of multiple sclerosis: implications for remyelination at lesions sites. Exp Neurol 250, 348–352

41. Yang, J., Yan, Y., Xia, Y., Kang, T., Li, X., Ciric, B., Xu, H., Rostami, A., and Zhang, G. X. (2014) Neurotrophin 3 transduction augments remyelinating and immunomodulatory capacity of neural stem cells. Mol Ther 22, 440–450

42. Gao, Z., Wen, Q., Xia, Y., Yang, J., Gao, P., Zhang, N., Li, H., and Zou, S. (2014) Osthole augments therapeutic efficiency of neural stem cells-based therapy in experimental autoimmune encephalomyelitis. J Pharmacol Sci 124, 54–65

43. Castelo-Branco, G., Stridh, P., Guerreiro-Cacais, A. O., Adzemovic, M. Z., Falcao, A. M., Marta, M., Berglund, R., Gillett, A., Hamza, K. H., Lassmann, H., Hermanson, O., and Jagodic, M. (2014) Acute treatment with valproic acid and l-thyroxine ameliorates clinical signs of experimental autoimmune encephalomyelitis and prevents brain pathology in DA rats. Neurobiol Dis 71, 220–233

44. Gao, Z., Nissen, J. C., Legakis, L., and Tsirka, S. E. (2015) Nicotine modulates neurogenesis in the central canal during experimental autoimmune encephalomyelitis. Neuroscience 297, 11–21

45. Theotokis, P., Kleopa, K. A., Touloumi, O., Lagoudaki, R., Lourbopoulos, A., Nousiopoulou, E., Kesidou, E., Poulatsidou, K. N., Dardiotis, E., Hadjigeorgiou, G., Karacostas, D., Cifuentes-Diaz, C., Irinopoulou, T., and Grigoriadis, N. (2015) Connexin43 and connexin47 alterations after neural precursor cells transplantation in experimental autoimmune encephalomyelitis. Glia 63, 1772–1783

46. Rafieemehr, H., Kheyrandish, M., and Soleimani, M. (2015) Neuroprotective Effects of Transplanted Mesenchymal Stromal Cells-derived Human Umbilical Cord Blood Neural Progenitor Cells in EAE. Iran J Allergy Asthma Immunol 14, 596–604

47. Gao, X., Hu, G., Deng, L., Fan, G., Yang, C., and Du, J. (2016) Transplantation of Neural Stem Cells Cotreated with Thyroid Hormone and GDNF Gene Induces Neuroprotection in Rats of Chronic Experimental Allergic Encephalomyelitis. Neural Plast 2016, 3081939

48. Rao, V. T., Khan, D., Jones, R. G., Nakamura, D. S., Kennedy, T. E., Cui, Q. L., Rone, M. B., Healy, L. M., Watson, R., Ghosh, S., and Antel, J. P. (2016) Potential Benefit of the Charge-Stabilized Nanostructure Saline RNS60 for Myelin Maintenance and Repair. Sci Rep 6, 30020

49. Covacu, R., and Brundin, L. (2019) Endogenous spinal cord stem cells in multiple sclerosis and its animal model. J Neuroimmunol 331, 4–10

50. Zhang, Y., Li, X., Ciric, B., Ma, C. G., Gran, B., Rostami, A., and Zhang, G. X. (2017) Effect of Fingolimod on Neural Stem Cells: A Novel Mechanism and Broadened Application for Neural Repair. Mol Ther 25, 401–415

51. Shirazi, H. A., Rasouli, J., Ciric, B., Wei, D., Rostami, A., and Zhang, G. X. (2017) 1,25-Dihydroxyvitamin D3 suppressed experimental autoimmune encephalomyelitis through both immunomodulation and oligodendrocyte maturation. Exp Mol Pathol 102, 515–521

52. Ghareghani, M., Zibara, K., Sadeghi, H., Dokoohaki, S., Sadeghi, H., Aryanpour, R., and Ghanbari, A. (2017) Fluvoxamine stimulates oligodendrogenesis of cultured neural stem cells and attenuates inflammation and demyelination in an animal model of multiple sclerosis. Sci Rep 7, 4923

53. Le Blanc, K., Rasmusson, I., Gotherstrom, C., Seidel, C., Sundberg, B., Sundin, M., Rosendahl, K., Tammik, C., and Ringden, O. (2004) Mesenchymal stem cells inhibit the expression of CD25 (interleukin-2 receptor) and CD38 on phytohaemagglutinin-activated lymphocytes. Scand J Immunol 60, 307–315

54. Glennie, S., Soeiro, I., Dyson, P. J., Lam, E. W., and Dazzi, F. (2005) Bone marrow mesenchymal stem cells induce division arrest anergy of activated T cells. Blood 105, 2821–2827

55. Groh, M. E., Maitra, B., Szekely, E., and Koc, O. N. (2005) Human mesenchymal stem cells require monocyte-mediated activation to suppress alloreactive T cells. Exp Hematol 33, 928–934

56. Ramasamy, R., Tong, C. K., Seow, H. F., Vidyadaran, S., and Dazzi, F. (2008) The immunosuppressive effects of human bone marrow-derived mesenchymal stem cells target T cell proliferation but not its effector function. Cell Immunol 251, 131–136

57. Kronsteiner, B., Wolbank, S., Peterbauer, A., Hackl, C., Redl, H., van Griensven, M., and Gabriel, C. (2011) Human mesenchymal stem cells from adipose tissue and amnion influence T-cells depending on stimulation method and presence of other immune cells. Stem Cells Dev 20, 2115–2126

58. Yan, F., Liu, O., Zhang, H., Zhou, Y., Zhou, D., Zhou, Z., He, Y., Tang, Z., and Wang, S. (2019) Human dental pulp stem cells regulate allogeneic NK cells’ function via induction of anti-inflammatory purinergic signalling in activated NK cells. Cell Prolif 52, e12595

59. Ishida, N., Ishiyama, K., Saeki, Y., Tanaka, Y., and Ohdan, H. (2019) Cotransplantation of preactivated mesenchymal stem cells improves intraportal engraftment of islets by inhibiting liver natural killer cells in mice. Am J Transplant

60. Qu, M., Cui, J., Zhu, J., Ma, Y., Yuan, X., Shi, J., Guo, D., and Li, C. (2015) Bone marrow-derived mesenchymal stem cells suppress NK cell recruitment and activation in PolyI:C-induced liver injury. Biochem Biophys Res Commun 466, 173–179

61. Lu, Y., Liu, J., Liu, Y., Qin, Y., Luo, Q., Wang, Q., and Duan, H. (2015) TLR4 plays a crucial role in MSC-induced inhibition of NK cell function. Biochem Biophys Res Commun 464, 541–547

62. Di Ianni, M., Del Papa, B., De Ioanni, M., Moretti, L., Bonifacio, E., Cecchini, D., Sportoletti, P., Falzetti, F., and Tabilio, A. (2008) Mesenchymal cells recruit and regulate T regulatory cells. Exp Hematol 36, 309–318

63. Casiraghi, F., Azzollini, N., Cassis, P., Imberti, B., Morigi, M., Cugini, D., Cavinato, R. A., Todeschini, M., Solini, S., Sonzogni, A., Perico, N., Remuzzi, G., and Noris, M. (2008) Pretransplant infusion of mesenchymal stem cells prolongs the survival of a semiallogeneic heart transplant through the generation of regulatory T cells. J Immunol 181, 3933–3946

64. Carrion, F., Nova, E., Ruiz, C., Diaz, F., Inostroza, C., Rojo, D., Monckeberg, G., and Figueroa, F. E. (2010) Autologous mesenchymal stem cell treatment increased T regulatory cells with no effect on disease activity in two systemic lupus erythematosus patients. Lupus 19, 317–322

65. Zhang, X., Ren, X., Li, G., Jiao, C., Zhang, L., Zhao, S., Wang, J., Han, Z. C., and Li, X. (2011) Mesenchymal stem cells ameliorate experimental autoimmune uveoretinitis by comprehensive modulation of systemic autoimmunity. Invest Ophthalmol Vis Sci 52, 3143–3152

66. Gonzalez, M. A., Gonzalez-Rey, E., Rico, L., Buscher, D., and Delgado, M. (2009) Treatment of experimental arthritis by inducing immune tolerance with human adipose-derived mesenchymal stem cells. Arthritis Rheum 60, 1006–1019

67. Bouffi, C., Bony, C., Jorgensen, C., and Noel, D. (2011) Skin fibroblasts are potent suppressors of inflammation in experimental arthritis. Ann Rheum Dis 70, 1671–1676

68. Cappellesso-Fleury, S., Puissant-Lubrano, B., Apoil, P. A., Titeux, M., Winterton, P., Casteilla, L., Bourin, P., and Blancher, A. (2010) Human fibroblasts share immunosuppressive properties with bone marrow mesenchymal stem cells. J Clin Immunol 30, 607–619

69. Zhang, X., Kiapour, N., Kapoor, S., Khan, T., Thamilarasan, M., Tao, Y., Cohen, S., Miller, R., Sobel, R. A., and Markovic-Plese, S. (2019) IL-11 Induces Encephalitogenic Th17 Cells in Multiple Sclerosis and Experimental Autoimmune Encephalomyelitis. J Immunol

70. de Oliveira Boldrini, V., Dos Santos Farias, A., and Degasperi, G. R. (2019) Deciphering targets of Th17 cells fate: From metabolism to nuclear receptors. Scand J Immunol, e12793

71. Shen, H., and Shi, L. Z. (2019) Metabolic regulation of TH17 cells. Mol Immunol 109, 81–87

72. Cho, J. J., Xu, Z., Parthasarathy, U., Drashansky, T. T., Helm, E. Y., Zuniga, A. N., Lorentsen, K. J., Mansouri, S., Cho, J. Y., Edelmann, M. J., Duong, D. M., Gehring, T., Seeholzer, T., Krappmann, D., Uddin, M. N., Califano, D., Wang, R. L., Jin, L., Li, H., Lv, D., Zhou, D., Zhou, L., and Avram, D. (2019) Hectd3 promotes pathogenic Th17 lineage through Stat3 activation and Malt1 signaling in neuroinflammation. Nat Commun 10, 701

73. Iannello, A., Rolla, S., Maglione, A., Ferrero, G., Bardina, V., Inaudi, I., De Mercanti, S., Novelli, F., D’Antuono, L., Cardaropoli, S., Todros, T., Turrini, M. V., Cordioli, C., Puorro, G., Marsili, A., Lanzillo, R., Brescia Morra, V., Cordero, F., De Bortoli, M., Durelli, L., Visconti, A., Cutrupi, S., and Clerico, M. (2018) Pregnancy Epigenetic Signature in T Helper 17 and T Regulatory Cells in Multiple Sclerosis. Front Immunol 9, 3075

74. Zhang, X., Kiapour, N., Kapoor, S., Merrill, J. R., Xia, Y., Ban, W., Cohen, S. M., Midkiff, B. R., Jewells, V., Shih, Y. I., and Markovic-Plese, S. (2018) IL-11 antagonist suppresses Th17 cell-mediated neuroinflammation and demyelination in a mouse model of relapsing-remitting multiple sclerosis. Clin Immunol 197, 45–53

75. Darlington, P. J., Stopnicki, B., Touil, T., Doucet, J. S., Fawaz, L., Roberts, M. E., Boivin, M. N., Arbour, N., Freedman, M. S., Atkins, H. L., and Bar-Or, A. (2018) Natural Killer Cells Regulate Th17 Cells After Autologous Hematopoietic Stem Cell Transplantation for Relapsing Remitting Multiple Sclerosis. Front Immunol 9, 834

76. Kurte, M., Luz-Crawford, P., Vega-Letter, A. M., Contreras, R. A., Tejedor, G., Elizondo-Vega, R., Martinez-Viola, L., Fernandez-O’Ryan, C., Figueroa, F. E., Jorgensen, C., Djouad, F., and Carrion, F. (2018) IL17/IL17RA as a Novel Signaling Axis Driving Mesenchymal Stem Cell Therapeutic Function in Experimental Autoimmune Encephalomyelitis. Front Immunol 9, 802

77. Sacramento, P. M., Monteiro, C., Dias, A. S. O., Kasahara, T. M., Ferreira, T. B., Hygino, J., Wing, A. C., Andrade, R. M., Rueda, F., Sales, M. C., Vasconcelos, C. C., and Bento, C. A. M. (2018) Serotonin decreases the production of Th1/Th17 cytokines and elevates the frequency of regulatory CD4(+) T-cell subsets in multiple sclerosis patients. Eur J Immunol 48, 1376–1388

78. van Langelaar, J., van der Vuurst de Vries, R. M., Janssen, M., Wierenga-Wolf, A. F., Spilt, I. M., Siepman, T. A., Dankers, W., Verjans, G., de Vries, H. E., Lubberts, E., Hintzen, R. Q., and van Luijn, M. M. (2018) T helper 17.1 cells associate with multiple sclerosis disease activity: perspectives for early intervention. Brain 141, 1334–1349

79. Zhang, Z., Xue, Z., Liu, Y., Liu, H., Guo, X., Li, Y., Yang, H., Zhang, L., Da, Y., Yao, Z., and Zhang, R. (2018) MicroRNA-181c promotes Th17 cell differentiation and mediates experimental autoimmune encephalomyelitis. Brain Behav Immun 70, 305–314

80. Kaushansky, N., Kaminitz, A., Allouche-Arnon, H., and Ben-Nun, A. (2019) Modulation of MS-like disease by a multi epitope protein is mediated by induction of CD11c(+)CD11b(+)Gr1(+) myeloid-derived dendritic cells. J Neuroimmunol 333, 476953

81. Xie, Z., Chen, J., Zheng, C., Wu, J., Cheng, Y., Zhu, S., Lin, C., Cao, Q., Zhu, J., and Jin, T. (2017) 1,25-dihydroxyvitamin D3 -induced dendritic cells suppress experimental autoimmune encephalomyelitis by increasing proportions of the regulatory lymphocytes and reducing T helper type 1 and type 17 cells. Immunology 152, 414–424

82. Buckland, M., Jago, C. B., Fazekasova, H., Scott, K., Tan, P. H., George, A. J., Lechler, R., and Lombardi, G. (2006) Aspirin-treated human DCs up-regulate ILT-3 and induce hyporesponsiveness and regulatory activity in responder T cells. Am J Transplant 6, 2046–2059

83. Ureta, G., Osorio, F., Morales, J., Rosemblatt, M., Bono, M. R., and Fierro, J. A. (2007) Generation of dendritic cells with regulatory properties. Transplant Proc 39, 633–637

84. Gaudreau, S., Guindi, C., Menard, M., Besin, G., Dupuis, G., and Amrani, A. (2007) Granulocyte-macrophage colony-stimulating factor prevents diabetes development in NOD mice by inducing tolerogenic dendritic cells that sustain the suppressive function of CD4+CD25+ regulatory T cells. J Immunol 179, 3638–3647

85. Zhang, X., Li, M., Lian, D., Zheng, X., Zhang, Z. X., Ichim, T. E., Xia, X., Huang, X., Vladau, C., Suzuki, M., Garcia, B., Jevnikar, A. M., and Min, W. P. (2008) Generation of therapeutic dendritic cells and regulatory T cells for preventing allogeneic cardiac graft rejection. Clin Immunol 127, 313–321

86. Cools, N., Van Tendeloo, V. F., Smits, E. L., Lenjou, M., Nijs, G., Van Bockstaele, D. R., Berneman, Z. N., and Ponsaerts, P. (2008) Immunosuppression induced by immature dendritic cells is mediated by TGF-beta/IL-10 double-positive CD4+ regulatory T cells. J Cell Mol Med 12, 690–700

87. Wolfle, S. J., Strebovsky, J., Bartz, H., Sahr, A., Arnold, C., Kaiser, C., Dalpke, A. H., and Heeg, K. (2011) PD-L1 expression on tolerogenic APCs is controlled by STAT-3. Eur J Immunol 41, 413–424

88. Tateosian, N. L., Reiteri, R. M., Amiano, N. O., Costa, M. J., Villalonga, X., Guerrieri, D., and Maffia, P. C. (2011) Neutrophil elastase treated dendritic cells promote the generation of CD4(+)FOXP3(+) regulatory T cells in vitro. Cell Immunol 269, 128–134

89. Volchenkov, R., Karlsen, M., Jonsson, R., and Appel, S. (2013) Type 1 regulatory T cells and regulatory B cells induced by tolerogenic dendritic cells. Scand J Immunol 77, 246–254

90. Farias, A. S., Spagnol, G. S., Bordeaux-Rego, P., Oliveira, C. O., Fontana, A. G., de Paula, R. F., Santos, M. P., Pradella, F., Moraes, A. S., Oliveira, E. C., Longhini, A. L., Rezende, A. C., Vaisberg, M. W., and Santos, L. M. (2013) Vitamin D3 induces IDO+ tolerogenic DCs and enhances Treg, reducing the severity of EAE. CNS Neurosci Ther 19, 269–277

91. Matsumoto, T., Hasegawa, H., Onishi, S., Ishizaki, J., Suemori, K., and Yasukawa, M. (2013) Protein kinase C inhibitor generates stable human tolerogenic dendritic cells. J Immunol 191, 2247–2257

92. Wu, Z. (2013) Antigen specific immunotherapy generates CD27(+) CD35(+) tolerogenic dendritic cells. Cell Immunol 283, 75–80

93. Kapina, M. A., Rubakova, E. I., Majorov, K. B., Logunova, N. N., and Apt, A. S. (2013) Capacity of lung stroma to educate dendritic cells inhibiting mycobacteria-specific T-cell response depends upon genetic susceptibility to tuberculosis. PLoS One 8, e72773

94. Pletinckx, K., and Lutz, M. B. (2014) Dendritic cells generated with Flt3L and exposed to apoptotic cells lack induction of T cell anergy and Foxp3(+) regulatory T cell conversion in vitro. Immunobiology 219, 230–240

95. Nessler, J., Benardais, K., Gudi, V., Hoffmann, A., Salinas Tejedor, L., Janssen, S., Prajeeth, C. K., Baumgartner, W., Kavelaars, A., Heijnen, C. J., van Velthoven, C., Hansmann, F., Skripuletz, T., and Stangel, M. (2013) Effects of murine and human bone marrow-derived mesenchymal stem cells on cuprizone induced demyelination. PLoS One 8, e69795

